# Interleukin-10 and Small Molecule SHIP1 Allosteric Regulators Trigger Anti-Inflammatory Effects Through SHIP1/STAT3 Complexes

**DOI:** 10.1101/2020.05.29.123943

**Authors:** Thomas C. Chamberlain, Sylvia T. Cheung, Jeff S.J. Yoon, Andrew Ming-Lum, Bernd R. Gardill, Soroush Shakibakho, Edis Dzananovic, Fuqiang Ban, Abrar Samiea, Kamaldeep Jawanda, John Priatel, Gerald Krystal, Christopher J. Ong, Artem Cherkasov, Raymond J. Andersen, Sean A. McKenna, Filip Van Petegem, Alice L-F Mui

**Author notes:** TCC, STC and JSJY contributed equally to this study. Lead Author: Address: Alice L-F Mui, Jack Bell Research Centre, 2660 Oak Street, Vancouver BC, V6H 3Z6, (Ext. 62242).

## Abstract

The anti-inflammatory actions of interleukin-10 (IL10) are thought to be mediated primarily by the STAT3 transcription factor, but pro-inflammatory cytokines such as interleukin-6 (IL6) also act through STAT3. We now report that IL10, but not IL6 signaling, induces formation of a complex between STAT3 and the inositol polyphosphate-5-phosphatase SHIP1 in macrophages. Both SHIP1 and STAT3 translocate to the nucleus in macrophages. Remarkably, sesquiterpenes of the Pelorol family we previously described as allosteric activators of SHIP1 phosphatase activity, could induce SHIP1/STAT3 complex formation in cells, and mimic the anti-inflammatory action of IL10 in a mouse model of colitis. Using crystallography and docking studies we identified a drug-binding pocket in SHIP1. Our studies reveal new mechanisms of action for both STAT3 and SHIP1, and provide a rationale for use of allosteric SHIP1-activating compounds which mimic the beneficial anti-inflammatory actions of IL10.

## INTRODUCTION

The prevalence of inflammatory bowel disease (IBD) in North America is 505/100,000 persons (Ulcerative Colitis) and 322/100,000 person (Crohn’s Disease) and increasing (Sairenji et al., 2017). Many factors contribute to the development of IBD, but genome wide association studies (Verstockt et al., 2018) and clinical data (Engelhardt and Grimbacher, 2014, Glocker et al., 2009, Glocker et al., 2011, Louis et al., 2009) show the anti-inflammatory actions of interleukin-10 (IL10) (Friedrich et al., 2019, Ouyang and O’Garra, 2019, Ouyang et al., 2011) are important in maintaining proper immune homeostasis. IL10 deficient mice develop colitis similar to human IBD (Kuhn et al., 1993, Shouval et al., 2014b) and the key target of IL10 is the macrophage (Friedrich et al., 2019, Shouval et al., 2014b, Zigmond et al., 2014). In humans, polymorphisms in the IL10 gene are associated with ulcerative colitis (Louis et al., 2009) and homozygous loss-of-function mutations in the IL10 receptor subunits result in early onset colitis (Engelhardt and Grimbacher, 2014, Glocker et al., 2009, Glocker et al., 2011). Therefore, understanding the mechanism by which IL10 exerts its action on target cells may provide insight into the development of therapeutics to treat inflammatory disease (Kumar et al., 2017).

IL10 maintains colon mucosal immune homeostasis mainly by inhibiting macrophage production of inflammatory mediators such as tumor necrosis factor α (TNFα) and IL1α elicited by inflammatory stimuli (Friedrich et al., 2019, Ouyang and O’Garra, 2019, Ouyang et al., 2011, Shouval et al., 2014a, Zigmond et al., 2014, Iyer and Cheng, 2012). In the classic model of IL10 receptor signaling, binding of IL10 to its receptor induces activation of the Jak1 and Tyk2 tyrosine kinase, tyrosine phosphorylation of the STAT3 transcription factor and expression of STAT3-regulated genes (Hutchins et al., 2013, Murray, 2006b, Murray, 2006a). STAT3 activation is widely thought to be sufficient to mediate all the anti-inflammatory actions of IL10 (El Kasmi et al., 2006, Murray, 2005, Murray, 2006b, Murray, 2006a, Weaver et al., 2007, Hutchins et al., 2012). In support of this, a myeloid specific STAT3^-/-^ mouse develops colitis (Takeda et al., 1999) much like an IL10^-/-^ mouse (Kuhn et al., 1993, Zigmond et al., 2014). However, STAT3 becomes tyrosine phosphorylated and activated by many stimuli including the pro-inflammatory cytokine interleukin-6 (IL6) (Garbers et al., 2015), so STAT3 activation must differ downstream of IL10 and IL6 signaling in order to mediate their opposing actions.

The SHIP1 phosphatidylinositol 3,4,5-trisphosphate 5-phosphatase is a cytoplasmic protein expressed predominantly in hematopoietic cells (Hibbs et al., 2018, Fernandes, 2013 #1400, Huber et al., 1999, Krystal, 2000, Pauls and Marshall, 2017). In response to extracellular signals, SHIP1 can be recruited to the cell membrane and one of its actions can be to turn off phosphoinositide 3-kinase (PI3K) signaling (Brown et al., 2010) by dephosphorylating the PI3K product PIP3 into PI(3,4)P_2_ (Fernandes et al., 2013, Huber et al., 1999, Krystal, 2000, Pauls and Marshall, 2017). We have shown that SHIP1 phosphatase activity is allosterically activated by its product PI(3,4)P_2_ and that small molecules of the pelorol family (ZPR-MN100 and ZPR-151) also allosterically enhance SHIP1 phosphatase activity (Meimetis et al., 2012, Ong et al., 2007). These data suggest that stimulating SHIP1 phosphatase activity with small molecule SHIP1 activators could be used to treat inflammatory diseases caused by inappropriately sustained PI3K production of PI(3,4)P_2_.

However, in addition to its enzymatic function in hydrolyzing PIP3, SHIP1 can also act as a docking protein for assembly of signaling complexes (Pauls and Marshall, 2017). We previously showed that IL10R signaling requires SHIP1 to inhibit TNFα translation (Chan et al., 2012) but whether SHIP1 and STAT3 worked independently or together was not determined. We now report that a SHIP1 protein containing point mutations, which inactivates its phosphatase activity, could still mediate the anti-inflammatory action of IL10, and that SHIP1 and STAT3 associate with each other in response to IL10. Furthermore, small molecule allosteric activators of SHIP1 can by themselves induce SHIP1/STAT3 complex formation and inhibit inflammation in a mouse model of colitis.

These data suggest that SHIP1 agonists can be used to elicit the beneficial anti-inflammatory action of IL10 by inducing a conformational change in SHIP1 that allows SHIP1/STAT3 complex formation.

Furthermore, disease indications in which loss of normal IL10 function has been implicated, are ones which might benefit most from use of SHIP1 agonists.

## RESULTS

### IL10 requires both SHIP1 and STAT3 to inhibit macrophage production of TNFα

A role for STAT3 in mediating IL10 inhibition of TNFα *in vivo* was first described by Takeda *et al* (Takeda et al., 1999). They found that LPS administration to mice with a myeloid specific knockdown of STAT3 produced more TNFα than wild-type mice, concluding endogenous IL10 is unable to counteract LPS signaling in the STAT3^-/-^ mice. However, closer examination of their data showed that while TNFα levels remain high post LPS administration in the IL10^-/-^ mice (Berg et al., 1995), TNFα levels drop in STAT3^-/-^ mice as endogenous levels of IL10 rise (Takeda *et al*, Fig 2B). This implies that a protein other than STAT3 might contribute to IL10 action. Our previous cell culture-based studies suggested that SHIP1 participates in IL10 action (Chan et al., 2012, Cheung et al., 2013, Samiea et al., 2020). We now looked *in vivo* at the ability of IL10 to inhibit LPS-induced inflammatory cytokine expression (**Figure 1A**) in SHIP1^+/+^ and SHIP1^-/-^ mice, and found IL10 inhibited TNFα in SHIP1^+/+^ but not SHIP1^-/-^ mice. We have previously shown that SHIP1 phosphatase activity is allosterically stimulated by its product PI(3,4)P_2_ (Ong et al., 2007). We synthesized a small molecule allosteric regulator, **ZPR-MN100** (previously called AQX-MN100 (Ong et al., 2007)) (**Figure 7A**) which binds to the same SHIP1 C2 domain as PI(3,4)P_2_ and increases SHIP1 functional activity (Ong et al., 2007). We found that ZPR-MN100 inhibited LPS-induced TNFα in SHIP1^+/+^ but not SHIP1^-/-^ mice, suggesting that these compounds can mimic anti-inflammatory properties of IL10 and is indeed specific for SHIP1 (**Figure 1B**).

**Figure 1.**
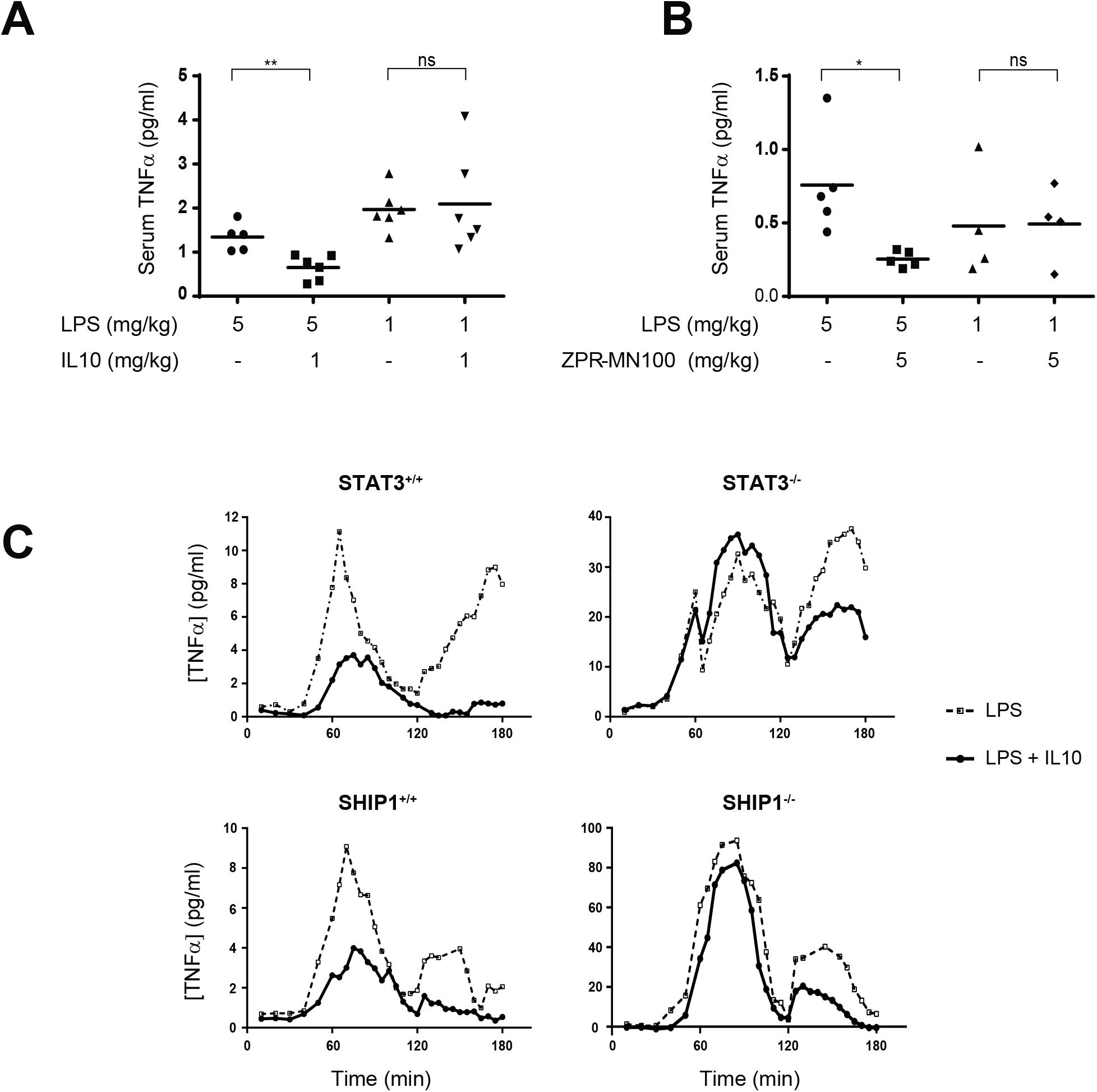
IL10 uses SHIP1 and STAT3 to inhibit macrophage activation. Serum TNFα in SHIP1^+/+^ or SHIP1^-/-^ mice injected intra-peritoneally with LPS, LPS + IL10 **(A)** or LPS + ZPR-MN100 **(B)** at the concentrations indicated for 1 hour. Data represent means of n≥4. * p<0.05, **p<0.01 when comparing to LPS alone stimulated mice, ns = not significant. **(C)** STAT3^+/+^, STAT3^-/-^, SHIP1^+/+^ and SHIP1^-/-^ bone marrow derived-macrophages (BMDMs) were stimulated with LPS (dotted line) or LPS + IL10 (solid line) over the course of 180 minutes in a continuous-flow apparatus. Fractions were collected every 5 minutes for measurement of TNFα levels. Data are representative of two independent experiments.

**Figure 2.**
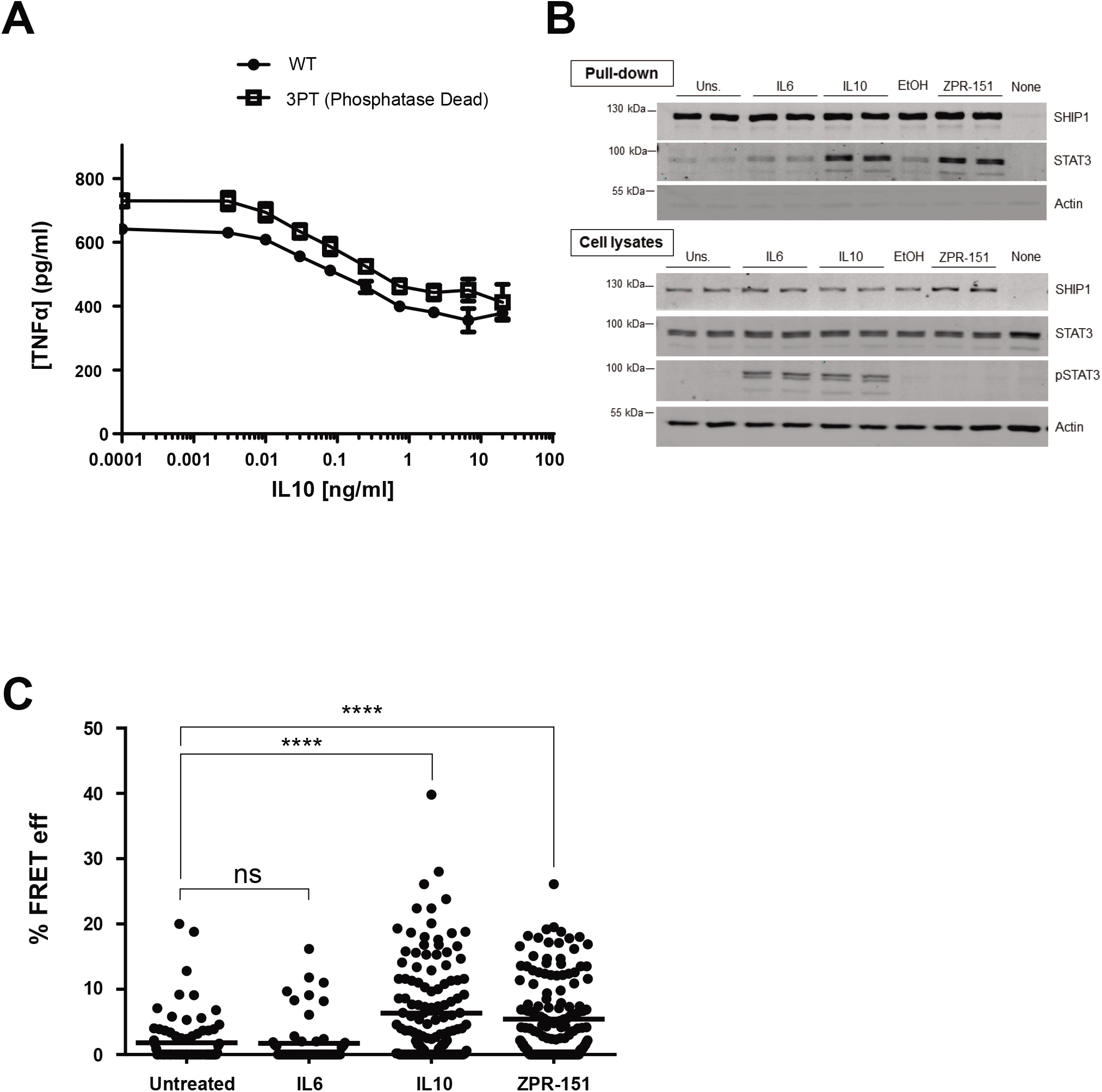
IL10 induces physical association of SHIP1 and STAT3. **(A)** J17 SHIP1^-/-^ cells expressing either His_6_-SHIP1 or His_6_-SHIP1 3PT were tested for their ability to be inhibited by IL10 in a LPS-stimulated TNFα production assay. **(B)** J17 His_6_-SHIP1 cells were stimulated with IL6, IL10 or ZPR-151 for 5 minutes. His_6_-SHIP1 was pulled down using Nickel beads and along with cell lysates probed with SHIP1, STAT3 and phospho-STAT3 antibodies. **(C)** Single cell FRET analysis of J17 SHIP1^-/-^ cells expressing FRET pair fusion constructs, Clover-SHIP1 and mRuby2-STAT3, ‘mock’ stimulated or stimulated with IL6, IL10, ZPR-151 for 1 minute. FRET efficiency was determined using the Acceptor Photobleaching method. Data represent % FRET efficiency of single cells from at least three independent experiments for each treatment (One-Way ANOVA with Tukey’s correction, **** p<0.0001).

SHIP1 and STAT3 could be acting independently or together in mediating IL10 action. To help distinguish between these two possibilities we used a continuous flow cell culture system that allows us to assess the kinetics of TNFα production in SHIP1 and STAT3 wild-type and knockout bone marrow derived macrophages (BMDM). LPS stimulates two peaks of TNFα expression, one at around 1 hour and another at 3 hours (**Figure 1C**). IL10 reduces TNFα levels in both SHIP1^+/+^ and STAT3^+/+^ cells, but is completely impaired in inhibiting the 1-hour peak in both STAT3^-/-^ and SHIP1^-/-^ cells, and partly impaired in inhibiting the 3-hour peak in both KO BMDM. The identical patterns of non-responsiveness suggest that SHIP1 and STAT3 cooperate.

### IL10 induces physical association of SHIP1 and STAT3 in macrophages

Our finding that SHIP1 and STAT3 work together is unprecedented. Both proteins reside in the cytoplasm in resting cells and are recruited to the cell membrane in response to extracellular stimuli but through distinct mechanisms. STAT3 functions mostly as a transcription factor (Matsuda et al., 2015) and SHIP1 is best known for its lipid phosphatase activity (Pauls and Marshall, 2017). However, SHIP1 can also act as a docking or adaptor protein for assembly of signaling complexes (Pauls and Marshall, 2017). Indeed, we found that a version of SHIP1 with minimal phosphatase activity (3PT) (An et al., 2005) can mediate the inhibitory effect of IL10 on LPS-stimulated TNFα production (**Figure 2A**), so we examined whether SHIP1 might serve an adaptor function in IL10 signaling and associate with STAT3 in response to IL10. **Figure 2B** shows that treatment of cells with IL10 resulted in co-precipitation of SHIP1 with STAT3. IL6 failed to induce STAT3 association with SHIP1, even though STAT3 becomes tyrosine phosphorylated to the same extent as in response to IL10. Remarkably, treatment of cells with the small molecule SHIP1 allosteric regulator **ZPR-151** (previously called 28·HCl (Meimetis et al., 2012)), a more water soluble derivative of ZPR-MN100 (**Figure 7A**), is sufficient to induce association of SHIP1 and STAT3 (**Figure 2B**). The ability of ZPR-151 to induce association of SHIP1 and STAT3 suggests the binding of ZPR-151 may induce a conformational change that can alter SHIP1’s association with other proteins. To see if the SHIP1/STAT3 interaction occurs in intact cells, we generated Clover-SHIP1 and mRuby2-STAT3 fusion protein constructs and transduced them into J17 SHIP1^-/-^ cells for FRET analysis. **Figure 2C** shows that stimulating Clover-SHIP1/mRuby2-STAT3 cells with IL10 or ZPR-151, but not IL6, increases the Clover-mRuby2 FRET signal suggesting SHIP1 and STAT3 interact *in vivo*.

Both SHIP1 and STAT3 have SH2 domains and both have been reported to become phosphorylated on tyrosine residues, so the complex formation might be mediated through a phospho-tyrosine/SH2 interaction. Since **Figure 2B** shows that STAT3 does not have to be phosphorylated to bind to SHIP1 (see ZPR-151 lane), we looked at whether tyrosine residues on SHIP1 might become phosphorylated to interact with the STAT3 SH2 domain. Four tyrosine residues in SHIP1 exist in the context of a STAT3 SH2 domain recognition sequence. We constructed SHIP1 mutants in which each of these residues are substituted with phenylalanine, expressed them in the J17 SHIP1^-/-^ macrophage cell line and tested the ability of IL10 to inhibit TNFα expression (**Figure 3A**) in these cells. Cells expressing the Y190F mutant behaved like a SHIP1^-/-^ (**Figure 3A**) cell. The Y190F mutant ability to interact with STAT3 was reduced 2-fold in response to IL10 and ZPR-151 (**Figure 3B** and **3C**), suggesting that part of the SHIP1 interaction with STAT3 required phosphorylation of SHIP1 Y190.

We also looked at the subcellular localization of SHIP1 and STAT3 in primary cells. Wild-type, SHIP1^-/-^ or STAT3^-/-^ peritoneal macrophages were stimulated with IL10 or ZPR-151 and stained with antibodies against SHIP1 or STAT3. **Figure 4A and Figure 4B** shows IL10 or ZPR-151 induced membrane association of both SHIP1 and STAT3 at 2 min in wild-type cells. SHIP1 does not translocate in STAT3^-/-^ cells, and STAT3 does not translocate in SHIP1^-/-^ cells (**Figure 4B**). At 20 min, both SHIP1 and STAT3 are found in the nucleus in wild-type cells, and translocation required cells to express both STAT3 and SHIP1. Thus, ZPR-151 can mimic IL10 in with respect to SHIP1 and STAT3 translocations.

**Figure 3.**
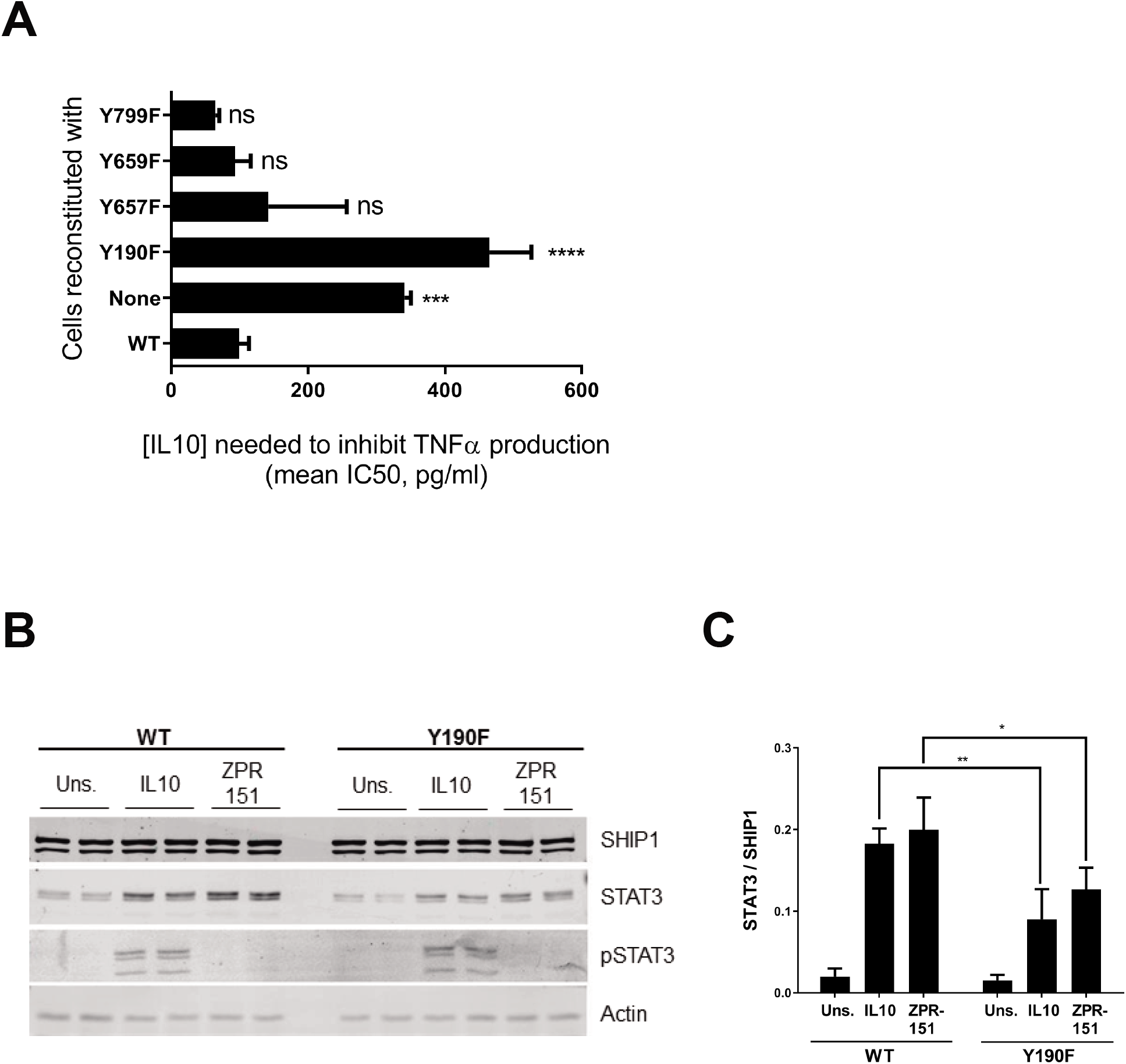
SHIP1 Y190 is involved in SHIP1 and STAT3 complex formation. **(A)** TNFα production of 1 ng/ml LPS + IL10 stimulated J17 SHIP1^-/-^ cells reconstituted with WT or mutant SHIP1 or vector (none) determined by ELISA from which IC50 values for IL10 were calculated (One-Way ANOVA with Dunnett’s correction **** p <0.0001). **(B)** Cells expressing either WT or Y190F SHIP1 were stimulated with IL10 or ZPR-151 for 5 minutes. His_6_-SHIP1 was pulled down using Nickel beads and along with cell lysates probed with SHIP1, STAT3 and phospho-STAT3 and actin antibodies. **(C)** The amount of STAT3 protein being pulled down with His_6_-SHIP1 (WT or Y190F) were quantified (Two-Way ANOVA with Sidak’s correction, ** p <0.01, * p <0.05).

**Figure 4.**
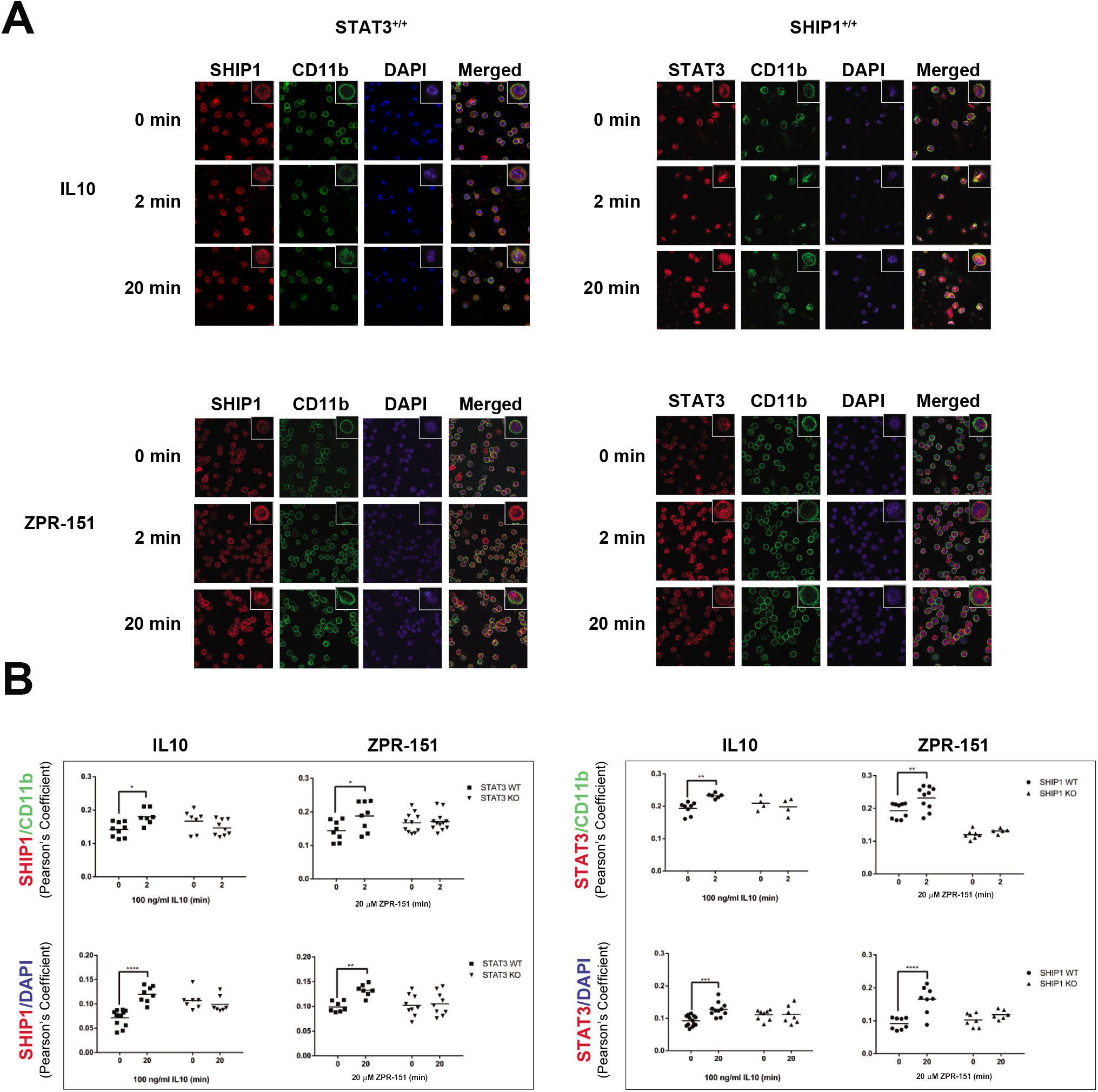
IL10 induces nuclear translocation of SHIP1 and STAT3. **(A)** STAT3^+/+^, and SHIP1^+/+^ perimacs were stimulated with IL10 or ZPR-151 for 2 or 20 minutes and stained with CD11b, SHIP1 and STAT3 antibodies and DAPI as indicated. Data for SHIP1^-/-^ and STAT3^-/-^ perimacs is shown in Figure S1. **(B)** Pearson’s coefficients were calculated to show the degree of overlap of SHIP1 or STAT3 with the membrane marker CD11b or DNA marker DAPI. Data represent Pearson’s coefficients for individual fields of cells from at least two independent experiments in each cell type (Two-Way ANOVA with Sidak’s correction, **** p<0.0001, *** p<0.001, ** p<0.01, * p<0.05).

### SHIP1 undergoes a conformational change upon allosteric regulator binding

To better understand the interaction of the small molecule allosteric regulators with SHIP1, we produced for X-ray crystallography, truncated SHIP1 proteins which contains the minimal region of SHIP1 needed for allosteric regulated phosphatase activity (**Figure 5**). Full length SHIP1 cannot be expressed at sufficient quantity for crystallography so we first defined the minimal region of SHIP1 needed for allosteric activation. We had previously shown the C2 domain binds the SHIP1 allosteric regulators (PI(3,4)P_2_, ZPR-MN100), and that the PH-R domain N-terminal to the phosphatase domain might be involved (Ong et al., 2007). So, we expressed full length SHIP1 (which could only be produced in mammalian 293T cells), **PPAC** (which contains the PH-R-phosphatase-C2 domains, and can be expressed in both 293T cells and *E. coli*), and **PAC1/PAC2** (which contain the phosphatase-C2 domains, and can be expressed in *E. coli*) proteins (**Figure 5A**). We examined their enzyme (phosphatase) kinetic properties and ability to become activated by ZPR-MN100. We found that 293T and *E. coli* derived PPAC had the same enzymatic properties (**Figure 5B**), and that all four proteins (full length SHIP1, 293T derived PPAC, *E. coli* derived PPAC, and PAC2) could be activated by ZPR-MN100 (**Figure 5C**).

**Figure 5.**
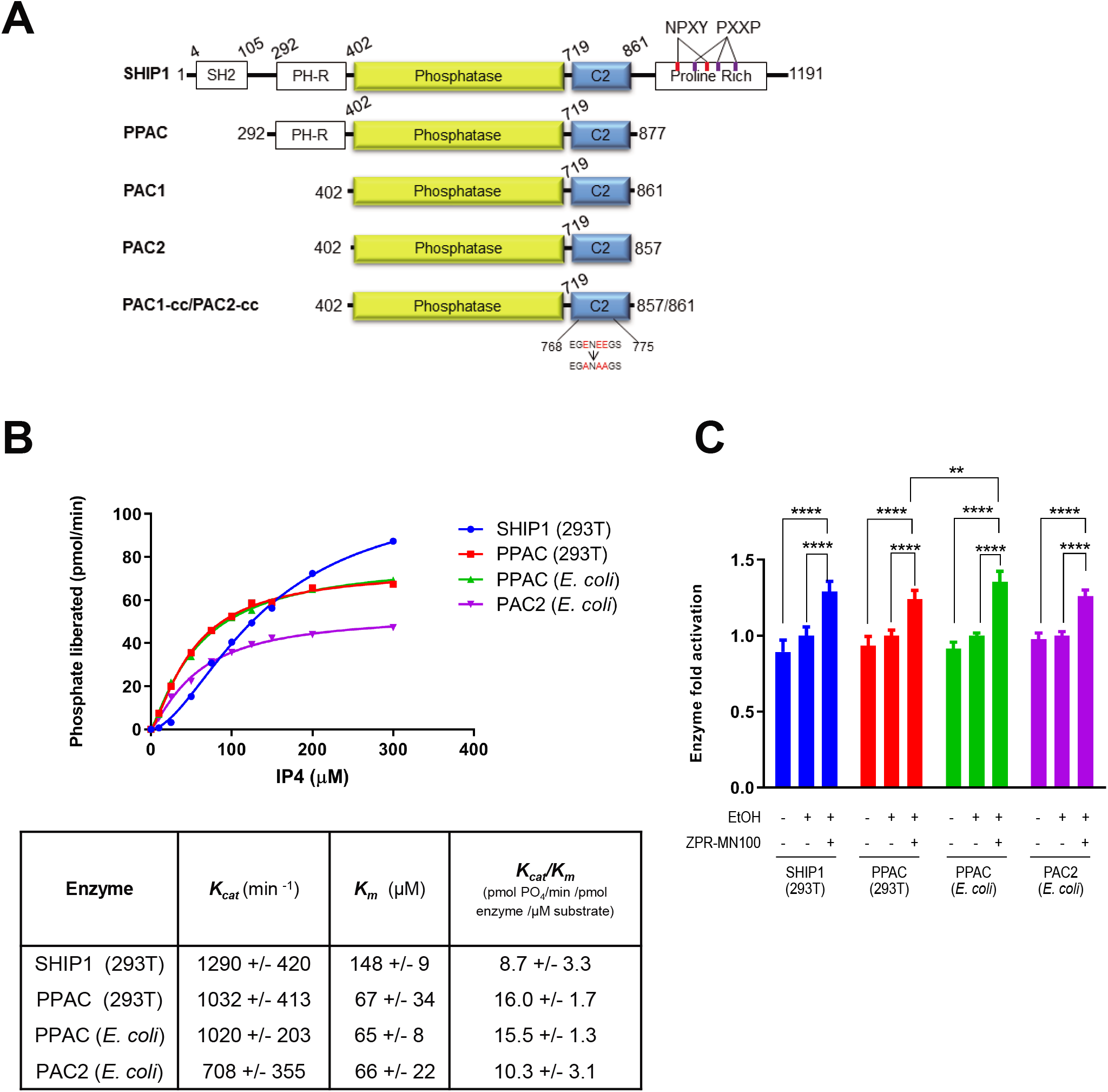
PPAC, PAC1 and PAC2 have similar enzymatic activity as full length SHIP1. **(A)** Schematic diagram of the different SHIP1 truncation constructs. PPAC consists of PH-R domain, phosphatase and C2 domain (residues aa 293-877) PAC1 and PAC2 consists of phosphatase and C2 domain (residues aa 402-861 and aa 402-857 respectively). PAC1-cc and PAC2-cc contain surface entropy reduction mutations in C2 domain (E770A, E772A, E773A). This cluster of residues were identified using the the SERp server (http://services.mbi.ucla.edu/SER/intro.php). **(B)** Enzyme catalytic initial velocities were determined at the indicated concentrations of IP_4_. K_cat_ and K_m_ values were calculated using GraphPad software **(C)** Ability of ZPR-MN100 to stimulate phosphatase activity in full length SHIP1, PPAC and PAC (Two-Way ANOVA with Tukey correction for multiple comparisons, **p<0.01, **** p<0.0001).

Only PAC1 and PAC2 proteins could be expressed in amounts needed for structural studies so these were produced and put through screens for conditions to produce crystals of sufficient quality for structural determination. This included making surface entropy reduced (Derewenda, 2004, Goldschmidt et al., 2007) versions called PAC1-cc and PAC2-cc where three glutamic acid residues in PAC1 and PAC2 were substituted with alanines. We solved the structure for several PAC1-cc and PAC2-cc crystals, and the data from a PAC2-cc crystal which diffracted at 1.6Å resolution is shown in **Figure 6A and Table S1**. We also used small angle X-ray scattering data (SAXS) to generate models of PAC1 with and without ZPR-MN100. The solution conformation of unliganded PAC1 determined by SAXS confirms the X-ray crystal structure. Furthermore, SAXS analysis showed that the binding of ZPR-MN100 to PAC1 results in a change in its overall conformation (**Figure 6B)**.

**Figure 6.**
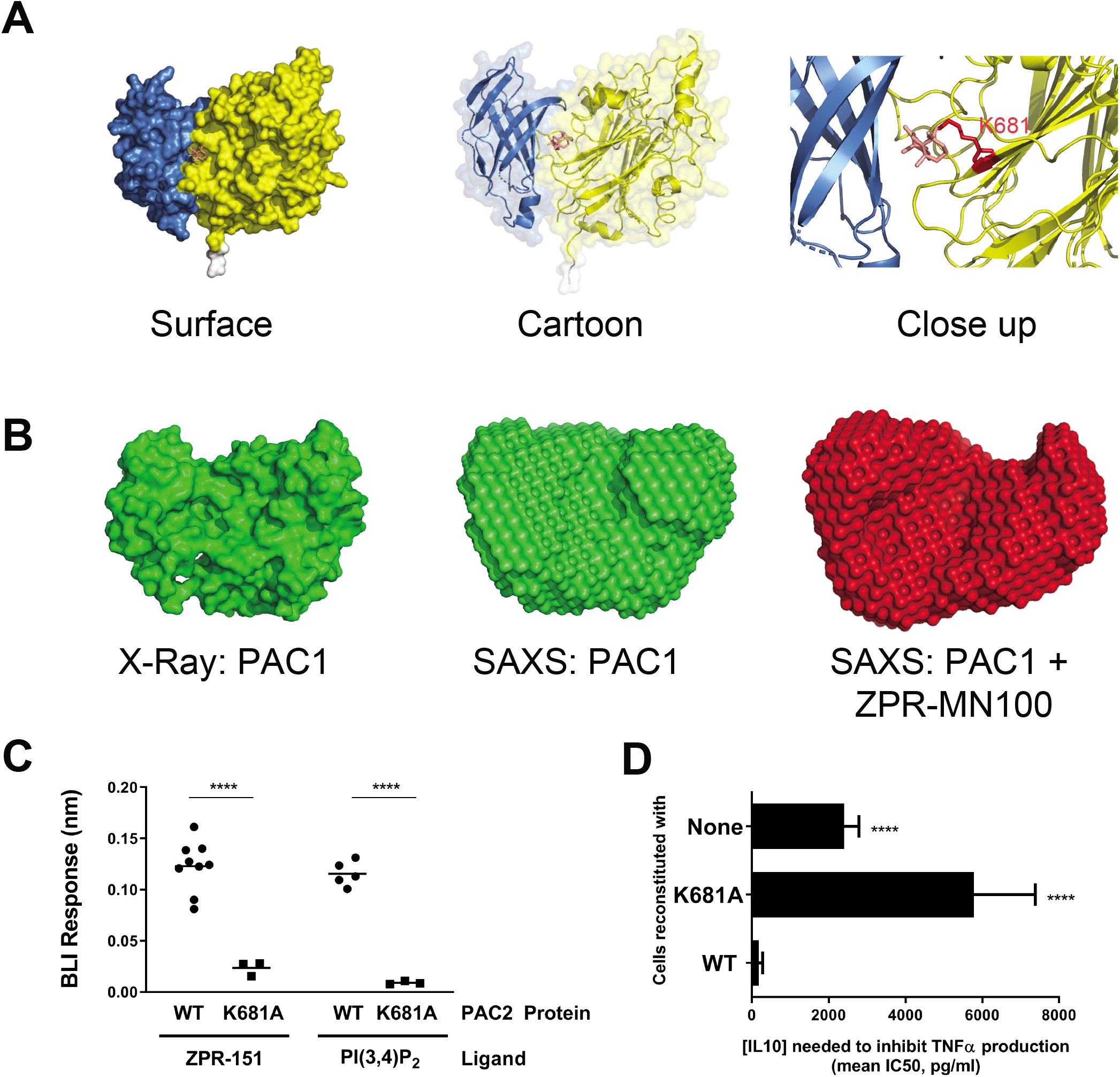
SHIP1 undergoes conformational change upon allosteric regulator binding. **(A)** Model of PAC2 based on crystallography data. The predicted binding pocket for the ZPR-MN100 (pink stick diagram) is located in the interface between C2 (blue) and phosphatase (yellow) domains. The binding pocket has amino acid residues K681 in close proximity with ZPR-MN100. **(B)** SAXS model of PAC1 in the absence of (apo-PAC1) and presence of liganded ZPR-MN100. **(C)** Bio-layer interferometry (BLI) data of PAC2 WT and K681A loaded sensors exposed to either 20 μM of ZPR-151 or PI(3,4)P_2_. **** p<0.0001 comparing WT PAC2 and K681A (Unpaired Student’s t-test) **(D)** TNFα production of 10 ng/ml LPS + IL10 stimulated cells reconstituted with WT or K681A SHIP1 or none (SHIP1 KO) determined by ELISA from which IC50 values for IL10 were calculated. **** p<0.0001 when comparing to cells reconstituted with WT SHIP1 (Unpaired Student’s t-test). See also Figure S2 and Table S1.

Using molecular modeling (Ban et al., 2018) we identified a potential binding pocket for ZPR-MN100/ZPR-151 in PAC2 (**Figure 6A)**. Residue K681 in this pocket is predicted to be involved in binding ZPR-MN100/ZPR-151 so we generated the K681A point mutant of PAC2 and tested the ability of wild-type and PAC2-K681A to bind ZPR-151 using Biolayer Interferometry (BLI). As shown in **Figure 6C** (and **Figure S2)**, substitution of K681A in the putative pocket impairs the ability of both ZPR-151 and PI(3,4)P_2_ to bind to PAC2. We then looked at the effect of the K681A substitution on the ability of full length SHIP1 to mediate IL10 inhibition of macrophages. **Figure 6D** shows that IL10 inhibited TNFα production efficiently in cells expressing wild-type but not K681A SHIP1.

In work done independently of the authors of this paper, Stenton *et al* described a molecule called AQX-1125 (structure in **Figure 7A**, later given the clinical trial name of Rosiptor) as a SHIP1 agonist (Stenton et al., 2013a, Stenton et al., 2013b). However, AQX-1125/Rosiptor has marginal SHIP1 phosphatase enhancing activity (Stenton et al., 2013b), and displayed different enzyme kinetics properties (Stenton et al., 2013b) than we observed with ZPR-MN100 (Ong et al., 2007). Stenton *et al* looked at binding of tritiated AQX-1125/Rosiptor to SHIP1 protein using scintillation proximity assay but it is difficult to assess the significance of the ~300 cpm signal they observed. So we directly compared the ability of PI(3,4)P_2_., ZPR-151 and AQX-1125/Rosiptor to bind to SHIP1 in our BLI assay (**Figure 7B and Figure S2**). We found AQX-1125/Rosiptor binds very poorly to SHIP1 as compared to ZPR-151 or SHIP1’s natural agonist PI(3,4)P_2_..

**Figure 7.**
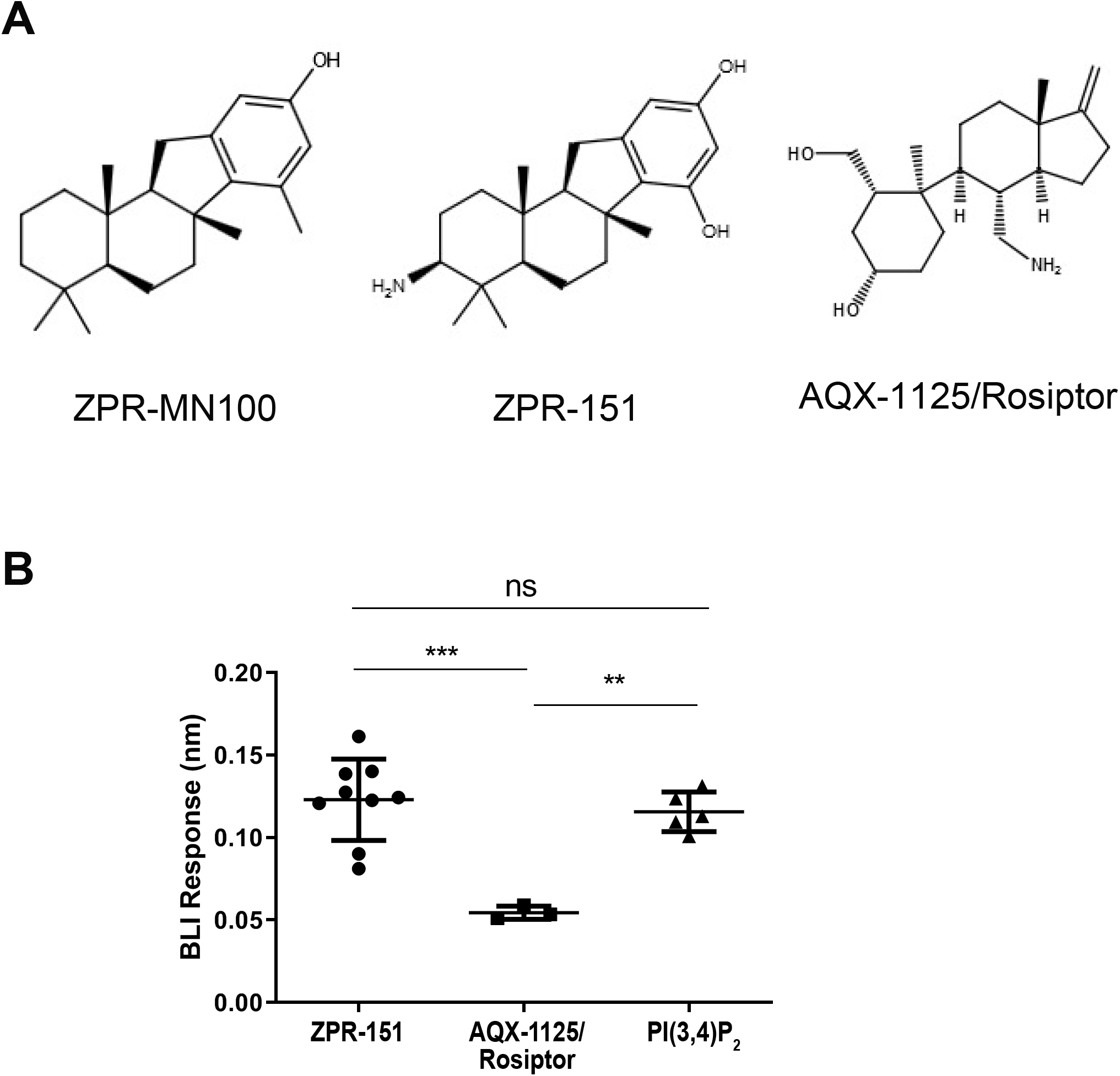
AQX-1125/Rosiptor binding to PAC2 is weak compared to ZPR-151 and PI(3,4)P_2_. **(A)** Structures of ZPR-MN100 and its derivative ZPR-151, and AQX-1125/Rosiptor. **(B)** Representative Bio-layer interferometry (BLI) curves for binding of 20 μM ZPR-151, AQX-1125 and PI(3,4)P_2_ to wild-type (WT) PAC2. Each data point indicates data from an independent biosensor (One-Way ANOVA with Tukey’s correction *** p <0.001, ** p<0.01). See also Figure S2.

### The small molecule SHIP1 allosteric regulator ZPR-MN100 alleviates inflammation in IL10^-/-^ colitis

To test whether small molecule SHIP1 allosteric regulators might be sufficient to mimic IL10’s anti-inflammatory action *in vivo*, we examined whether ZPR-MN100 could reduce inflammation in the IL10 knock-out mouse model of colitis (Keubler et al., 2015). IL10 knock-out mice develop colitis when colonized with normal gut flora because IL10 is needed to temper the host immune response to intestinal commensal bacteria (Keubler et al., 2015, Kuhn et al., 1993). We initiated colitis in IL10^-/-^ mice by inoculating them with the freshly isolated colon contents of normal, specific pathogen free mice and allowed inflammation to develop for 6 weeks (Sydora et al., 2003). Mice were then treated for 3 weeks with vehicle, 2 mg/kg ZPR-MN100, or 0.4 mg/kg dexamethasone (anti-inflammatory steroidal drug used as positive control) prior to colon tissue collection for analyses. Hematoxylin and eosin stained sections were prepared from the proximal, mid and distal colons of mice, as well as from mice not inoculated with flora (no colitis group) (**Figure 8A**). Two investigators blinded to the treatment groups scored the sections based on submucosal edema, immune cell infiltration, presence of goblet cells and epithelial integrity (**Figure 8B**). In the three groups where colitis was induced, the dexamethasone and ZPR-MN100 groups had significantly lower pathology scores than the vehicle group (**Figure 8B**). RNA was prepared from the colons of all four groups for analysis of IL17 and CCL2 expression, inflammatory mediators elevated in colitis (Lee et al., 2007). As shown in **Figure 8C**, both ZPR-MN100 and dexamethasone treatment significantly reduced the levels of IL17 and CCL2 mRNA. These data indicate that ZPR-MN100 treatment can reduce inflammation in colitis resulting from the loss of IL10.

## DISCUSSION

IL6 and IL10 have opposing pro- and anti-inflammatory actions respectively on macrophages (Garbers et al., 2015, Yasukawa et al., 2003) but both cytokines stimulate tyrosine phosphorylation of STAT3 Y705 in cells. We found that IL10 but not IL6 induced association of STAT3 with SHIP1, and suggest this difference may contribute to why STAT3 can mediate pro- and anti-inflammatory responses downstream of both cytokines. Perhaps IL10-induced SHIP1/STAT3 signaling support anti-inflammatory responses while IL6-induced STAT3/STAT3 dimers support pro-inflammatory responses. Yasukawa *et al*’s study of SOCS3 knockout cells suggested that the duration of STAT3 activation in macrophage cells can underlie the opposite biological effects of IL10 and IL6 (Yasukawa et al., 2003). Our data are compatible with theirs since STAT3 activation can also be prolonged by its association with SHIP1.

Our SHIP1 pulldown studies used an N-terminal His_6_-tagged SHIP1 construct transduced into a SHIP^-/-^ cell line. We were unable to co-precipitate SHIP1 and STAT3 with any other SHIP1 antibody (which are all directed to other regions of SHIP1). This suggests the N-terminus of SHIP1 is accessible in the SHIP1/STAT3 complex. Caveats of using cell lines transduced with His_6_-tagged SHIP1 include possible expression of SHIP1 to higher levels than seen in wild-type cells. However, as shown in **Figure S3**, the level of His_6_-tagged SHIP1 is similar to that of SHIP1 in SHIP^+/+^ cells. Another caveat is that associations seen in an *in vitro* pull-down experiment might not occur in intact cells. However, we were able to use FRET to show that Clover tagged SHIP1 and mRuby2 tagged STAT3 associate in cells in response to IL10 or ZPR-151 (**Figure 2C**). Future studies involve generation of antibodies to the N-terminus of SHIP1 in order to study endogenous SHIP1. We are interested in how IL10R signaling leads to SHIP1 working with STAT3 while IL6 receptor does not. We will also study whether a SHIP1/STAT3 complex induces expression of anti-inflammatory genes, or the presence of SHIP1 in the nucleus interferes with expression of IL6-induced, STAT3-regulated genes.

We previously showed that small molecule SHIP1 agonists have anti-inflammatory actions *in vitro* (Meimetis et al., 2012, Ong et al., 2007) and ascribed these actions to the stimulation of SHIP1’s phosphatase to dephosphorylate the PI3K product PIP3 into PI(3,4)P_2_ (Fernandes et al., 2013, Huber et al., 1999, Krystal, 2000, Pauls and Marshall, 2017). However, our current data demonstrate a SHIP1 protein with non-detectable phosphatase activity is sufficient to mediate the anti-inflammatory effect of IL10, so the adaptor function of SHIP1 can by itself support IL10 action. Our SAXS analyses suggest that the binding of SHIP1 agonists to SHIP1 causes a conformational change in SHIP1. This conformational change may allow SHIP1 to interact with STAT3 and the complex of SHIP1/STAT3 to translocate to the nucleus. We have solved the structure of the minimal domain (PAC1/2) of SHIP1 required to mediate the allosteric action of SHIP1 agonists, and identified a drug binding pocket through molecular docking analysis. Mutation of a residue predicted to be involved in binding to ZPR-151 abolished ZPR-151 binding and the ability of SHIP1 to mediate IL10 inhibition of TNFα expression in macrophages.

We found that SHIP1 Y190 contributes to the ability of SHIP1 to associate with STAT3. The Y190F mutant’s ability to interact with STAT3 was reduced 2-fold as compared to wild-type SHIP1 (**Figure 3B**). However, SHIP1 Y190F is completely impaired in its ability to support IL10 inhibition of TNFα (**Figure 3A**). One interpretation is the partial SHIP1/STAT3 complex inhibition is physiologically significant because inhibition of TNFα is completely abolished. Alternatively, the SHIP1/STAT3 complex formation is only one function of the Y190. Remarkably, the SHIP1 agonist ZPR-151 could by itself induce formation of a SHIP1/STAT3 complex. The addition of ZPR-151 to perimacs could also induce translocation of SHIP1 and STAT3 to the nucleus. Together these data suggest that the action of SHIP1 agonists includes both their ability to stimulate SHIP1 phosphatase activity and to induce the association of SHIP1 with STAT3.

Treatment of macrophages with ZPR-151/ZPR-MN100 was sufficient to elicit the anti-inflammatory effects similar to that of IL10 *in vitro* in this and our previous studies (Chan et al., 2012, Cheung et al., 2013, Ong et al., 2007). We thus tested ZPR-MN100 in a mouse model of colitis since the beneficial anti-inflammatory action of IL10 in colitis is through IL10 action on macrophages (Friedrich et al., 2019, Ouyang and O’Garra, 2019, Shouval et al., 2014b, Zigmond et al., 2014). We found that ZPR-MN100 was as effective as dexamethasone in reducing histological and molecular markers of colon inflammation. Medzhitov’s group recently reported IL10 stimulation of mitophagy and inactivation of the inflammasome as part of its protective effect in colitis, and that this involved STAT3-dependent upregulation of the DDIT4 protein (Ip et al., 2017). We confirmed IL10 upregulation of DDIT4 in macrophages requires both STAT3 and SHIP1; furthermore, ZPR-151 was by itself able to induce DDIT4 expression (manuscript in preparation).

A small molecule SHIP1 allosteric regulator (AQX-1125/Rosiptor) (Stenton et al., 2013a, Stenton et al., 2013b) developed independently of the authors of this manuscript was recently tested in clinical trials for relief of urinary bladder pain experienced by interstitial cystitis (IC) patients (Nickel et al., 2016). IC reportedly was chosen for the disease indication because: AQX-1125/Rosiptor accumulates in the urinary bladder (Stenton et al., 2013b), two papers implicated PI3K-dependent inflammation in IC (Liang et al., 2016, Qiao et al., 2014), and preliminary phase 2 trials seemed promising (Nickel et al., 2016). However, the phase 3 trial failed to show efficacy for AQX-1125/Rosiptor (Nickel et al., 2019). There are many reasons for small molecule drugs to fail during the drug development process. However, we note that neither IL10 nor SHIP1 has been implicated in the physiology/pathophysiology of IC (Nickel et al., 2019). Furthermore, we found that AQX-1125/Rosiptor has very weak binding to SHIP1, consistent with Stenton’s *et al* finding that AQX-1125 has very weak SHIP1 phosphatase activating ability (Stenton et al., 2013b).

We suggest that disease indications for which small molecule SHIP1 allosteric regulators are developed should be ones in which IL10) (Friedrich et al., 2019, Ouyang and O’Garra, 2019, Ouyang et al., 2011) (or other physiological regulators of SHIP1) (Hibbs et al., 2018, Chan et al., 2012, Cheung et al., 2013, Dobranowski and Sly, 2018, Pauls and Marshall, 2017) has been shown to play a beneficial role. These small molecules should also have similar binding properties to SHIP1 as its natural ligand PI(3,4)P_2_. By these criteria, small molecule SHIP1 agonists such as those of the Pelorol family should be explored for the treatment of human inflammatory bowel disease.

## ACKNOWLEDGEMENTS

We gratefully acknowledge Dr. Laura Sly for her critical review of the manuscript, and Dr. Nada Lallous for her advice for the BLI studies. Funding sources include: Canadian Institutes for Health Research (CIHR) (MOP-84539 to ALM), the Canadian Cancer Society Research Institute (Canadian Cancer Society grant # 017289 to ZPR), and an NSERC Discovery grant to SAM. STC and AML held CIHR Doctoral Research and Michael Smith Foundation for Health Research (MSFHR) awards. We thank the support staff at the Advanced Photon Source (Chicago) GM/CA-CAT beamline 23-ID-D, the Stanford Synchrotron Radiation Lightsource (Menlo Park, USA), and at the Canadian Light Source (Saskatoon, SK, Canada), which is supported by the Natural Sciences and Engineering Research Council of Canada, the National Research Council Canada, the Canadian Institutes of Health Research (CIHR), the Province of Saskatchewan, Western Economic Diversification Canada, and the University of Saskatchewan.

## AUTHOR CONTRIBUTIONS

Contribution: TCC, STC, JSJY, and ALM designed and performed research, analyzed data and wrote the manuscript. SS, AS, DS, SAM and KJ performed experiment(s) and analyzed data. BRG collected and analyzed crystallographic data. GK, CJO, JP, ZPR, SAM and FVP provided reagents and assisted with manuscript preparation.

Conflict of interest disclosure: ALM, RA, CO, and GK were scientific founders of Aquinox Pharmaceuticals but have not been associated with, or receive compensation from the company since 2010.

## CONTACT FOR REAGENT AND RESOURCE SHARING

Further information and requests for resources and reagents should be directed to and will be fulfilled by the Lead Contact, Alice Mui.

## EXPERIMENTAL MODEL AND SUBJECT DETAILS

### Mouse colonies

BALB/c mice wild type (^+/+^) or SHIP1 knockout (^-/-^) mice were provided by Dr. Gerald Krystal (BC Cancer Research Centre, Vancouver, BC). The generation of STAT3^-/-^ mice started with crossing C57BL/6 STAT3 flox/flox mice (Dr. Shizuo Akira, Hyogo College of Medicine, Nishinomiya, Japan) with C57BL/6 LysMcre mice (Jackson Laboratory). Their offspring were then crossed with homozygous STAT3 flox/flox mice to produce to generate both STAT3 flox/flox /LysMCre^+/-^ mice (referred to be STAT3^-/-^ mice) and STAT3 flox/flox mice (STAT3^+/+^ mice) in the same litters. All mice were maintained in accordance with the ethic protocols approved by the University of British Columbia Animal Care Committee.

### Constructs

The mammalian (lentiviral) expression plasmids of SHIP1 in FUGWBW were generated using Gateway LR reactions from pENTR1A (Invitrogen, Burlington, ON) constructs. A pENTR1A-His_6_-SHIP1 WT (SHIP1 Uniprot ID Q9ES52) plasmid was used as the template for standard primer based, site-directed mutagenesis to generate the K681A, Y190F, Y799F, Y659F and Y657F mutants. The phosphatase disrupted SHIP1 construct (P671A, D675A, and R676G in the phosphatase domain) was kindly provided by Dr. KS Ravichandran (University of Virginia). The constructs were confirmed by DNA sequencing. Subsequently, a Gateway LR reaction was performed between pENTR1A construct and FUGWBW (FUGW in which the green fluorescent protein was replaced by the Gateway cassette, and a blasticidin S resistance gene expression cassette was inserted downstream of the Gateway cassette (Peacock et al., 2009). Success of the LR reaction was confirmed by restriction enzyme digest. For Clover/mRuby2 based FRET experiments (Lam et al., 2012), pENTR1A Clover-SHIP1 was constructed by inserting a Clover fragment from pcDNA3 Clover (Addgene) to the N-terminal of SHIP1 in pENTR1A-His_6_-SHIP1 WT, replacing the His_6_.pENTR1A STAT3-mRuby2 was constructed by cloning murine STAT3 (Uniprot ID P42227) into pENTR1A followed by insertion of a mRuby2 fragment from pcDNA3 mRuby2 (Addgene) to the N-terminus of STAT3. Constructs were confirmed by sequencing and transferred to FUGWBW as above.

Bacteria expression vectors to produce recombinant proteins for crystallography and biolayer interferometry were generated by ligase-independent cloning (LIC) methodology in the LIC-HMT vector (Van Petegem et al., 2004). This plasmid contains an N-terminal tag composed of His_6_ and maltose binding protein (MBP), followed by a TEV protease cleavage site (abbreviated as the HMT-tag). The PCR product was purified and treated with T4 DNA polymerase (LIC-quality) (Novagen, Madison, WI) in the presence of dCTP only. The LIC-HMT vector was digested with *SspI* and the linearized plasmid was treated with T4 DNA polymerase in the presence of dGTP only. Equal volumes of insert and vector were mixed and incubated at room temperature for 10 minutes, followed by transformation into chemical competent *E. coli DH5α* cells using the standard heat shock protocol, and selection on kanamycin-containing LB agar plates. To generate different PAC2 mutants, standard site-directed mutagenesis was employed. Identities of all plasmids were confirmed by DNA sequencing.

### Cell lines

J16 and J17 cell lines derived from SHIP1^+/+^ and ^-/-^ BMDM respectively were previously described (Ming-Lum et al., 2012) and cultured in Mac media (IMDM supplemented with 10% (v/v) FCS, 10 μM β-mercaptoethanol, 150 μM monothioglycolate and 1 mM L-glutamine). J17 cells expressing wild type and mutant His_6_-SHIP1, Clover-SHIP1 or mRuby2-STAT3 constructs were generated by lentivirus mediated gene transfer as described (Cheung et al., 2013). Transduced cells were selected with 5 μg/ml blasticidin. Clover-SHIP1 and mRuby2-STAT3 cells were further subjected to fluorescent activated cell sorting to select the brightest cells on a FACS Aria II cytometer.

### Isolation of mouse peritoneal macrophages

Primary peritoneal macrophages (perimacs) were isolated from mice by peritoneal lavage with 3 ml of sterile Phosphate Buffered Saline (PBS) (Thermo Fisher Scientific, Nepean, ON). Perimacs were collected and transferred to Mac media.

### Production of bone marrow-derived macrophages

Bone marrow-derived macrophages (BMDMs) were generated by first collecting femurs and tibias from mice, and then flushing out the bone marrow through a 26-G needle. Extracted cells were plated, in Mac media supplemented with 5 ng/ml each of CSF-1 and GM-CSF (Stem Cell Technologies, Vancouver, BC), on a 10-cm tissue culture plate for 2 hours at 37°C. Non-adherent cells were collected and replated at 9×10^6^ cells per 10-cm tissue culture plate. Cells were then cultured in the presence of CSF-1 and GM-CSF. Differentiated BMDMs were used after 7 to 8 days. All cells were maintained in a 37°C, 5% CO_2_, 95% humidity incubator.

## METHOD DETAILS

### Continuous Flow Cultures

The continuous flow apparatus facilitates constant stimulation and removal of cell supernatants to determine kinetic profiles of cytokine production over time. BMDMs were seeded at 3×10^5^ cells per well in a 24-well tissue culture plate that had been coated with poly-L-lysine (Thermo Fisher Scientific, Nepean, ON) and rinsed with PBS. After overnight incubation, culture media was removed and Leibovitz’s L-15 (L-15) media (Invitrogen, Burlington, ON) supplemented with 3% FCS, 10 μM β-mercaptoethanol and 150 μM monothioglycolate was added. Cells were allowed to equilibrate in L-15 media for 1 hour before being placed in the continuous flow apparatus. Stimulation solution was made in the same media equilibrated at 37°C, and was passed through a modified inlet fitted to the well by a syringe pump (New Era Syringe Pumps Inc., Farmingdale, NY). A flow rate of 150 μl per minute was used. At the same time, cell supernatants were removed from the well at the same flow rate, and fractions were collected at 5-minute intervals over the course of 3 hours. These fractions were analyzed for secreted TNFα levels by ELISA.

### Real-time quantitative PCR

Total RNA was extracted using Trizol reagent (Invitrogen, Burlington, ON) according to manufacturer’s instructions. About 2-5 μg of RNA were treated with DNAseI (Roche Diagnostics, Laval, QC) according to the product manual. For mRNA expression analysis, 120 ng of RNA were used in the Transcriptor First Strand cDNA synthesis kit (Roche Diagnostics, Laval, QC), and 0.1 μl to 0.2 μl of cDNA generated were analyzed by SYBR Green-based real time PCR (real time-PCR) (Roche Diagnostics, Laval, QC) using 300 nM of gene-specific primers. Expression levels of mRNA were measured with the StepOne Plus RT-PCR system (Applied Biosystems, Burlington, ON), and the comparative Ct method was used to quantify mRNA levels using GAPDH as the normalization control.

### Measurement of TNFα production

Cells were seeded at 50 x 10^4^ cells per well in a 96-well tissue culture plate and allowed to adhere overnight. Media was changed the next day 1 hour prior to stimulation. Cells were stimulated with 1 or 10 ng/ml LPS +/-various concentrations of IL10 for 1 hour. Supernatant was collected and secreted TNFα protein levels were measured using a BD OptEIA Mouse TNFα Enzyme-Linked Immunosorbent Assay (ELISA) kit (BD Biosciences, Mississauga, ON). Triplicates wells were used for each stimulation condition. IC50 values were calculated from three independent experiments and differences in IL10 IC50 values from cells expressing SHIP1 mutants vs SHIP1 WT protein were analyzed by one-way ANOVA.

### *In vitro* phosphatase assay

The phosphatase assay was performed in 96-well microtiter plates with 10 ng of enzyme/well in a total volume of 25 μL in 20 mM Tris-HCl, pH 7.4, 150 mM NaCl, 0.05% Tween-20, 10 mM MgCl2 as described (Ong et al., 2007). Enzyme was incubated with or ZPR-MN100 (dissolved in ethanol) for 10 minutes at 23°C, before the addition of 50 μM of inositol-1,3,4,5-tetrakisphosphate (IP_4_) (Echelon Bioscience Inc., Salt Lake City, Utah). The reaction was allowed to proceed for 10 minutes at 37°C and the amount of inorganic phosphate released was assessed by the addition of Malachite Green reagent and absorbance measurement at 650 nm. For enzyme kinetic determination, different concentrations of IP_4_ (0 – 300 μM) were used and the reactions were stopped at different time points. Initial velocities were calculated, and K_cat_ and K_M_ were determined using GraphPad 6 software.

### *In vitro* pull down assay

J17 His_6_-SHIP1 and Y190F cells were seeded at 2 x 10^6^ cells per well in a 6-well plate. After overnight incubation, fresh cell media was added for 30 minutes before stimulation with 100 ng/ml IL10, IL6 or 20 μM ZPR-151 for 5 minutes. Cells were lysed with Protein Solubilization Buffer (PSB, 50 mM Hepes, pH 7.5, 100 nM NaF, 10 mM Na Pyrophosphate, 2 mM NaVO4, 2 mM Na Molybdate, 2 mM EDTA) containing 1% octylglucoside, 0.01 M imidazole, and protease inhibitor cocktail (Roche Diagnostics, Laval, QC) for 30 minutes at 4°C and centrifuged at 10000 rpm for 15 minutes. EDTA resistant Ni beads (Roche Diagnostics, Laval, QC) were added to supernatants and the mixture incubated at 4°C for 1 hour before spinning down and washing of beads three times with 0.1% octylglucoside wash buffer in PSB. Bead samples and starting material cell lysates were separated on a 7.5 % SDS-PAGE gel.

### Immunoblotting

Protein lysates were separated on SDS-PAGE gels and transferred onto polyvinylidene fluoride (PVDF) membrane (Millipore, Etobicoke, ON). Membranes were blocked and probed, where appropriate, with the following primary antibodies overnight: SHIP1 (P1C1) (Santa Cruz Biotechnology), pSHIP1 (Y190) (Kinexus), STAT3 (9D8) (ThermoFisher Scientific), pSTAT3 (Y705) (Thermo Fisher Scientific), STAT1 (BD Transduction Laboratories), pSTAT1 (Y701) (Upstate Biotechnology), GAPDH and actin (Sigma-Aldrich). Membranes were developed with either Alexa Fluor^®^ 660 anti-mouse IgG or Alexa Fluor^®^ 680 anti-rabbit IgG antibodies (Thermo Fisher Scientific) and imaged using a LI-COR Odyssey Imager.

### Acceptor Photobleaching FRET analysis

J17 SHIP1^-/-^cells expressing Clover-SHIP1 and/or mRuby2-STAT3 were seeded at 50 x 10^4^ cells per well in 8-well Ibidi μ-Slides (Ibidi GmbH, Martinsried, Germany). After overnight incubation, cells were serum-starved with Mac media containing 1% serum for 3 hours before media replacement with Leibovitz’s (L-15) media (Invitrogen, Burlington, ON) supplemented with 1% serum, 10 μM β-mercaptoethanol, 150 μM monothioglycolate and 1 mM L-glutamine for confocal microscopy imaging. Cells were imaged on a Leica SP8X on DMi8 confocal microscope system with a 63X/1.3 Gly HC PL APO CS2 objective using a white light laser line with 488 nm for donor excitement and 555 nm for acceptor excitement. Photobleaching FRET analysis was performed by measuring Clover-SHIP1 donor fluorescence intensity before and after bleaching the acceptor, mRuby2-STAT3, within a field of cells ‘region of interest’ (ROI), at 100% laser intensity for 60 frames. Acceptor photobleaching was performed first in resting cells then at 1 minute following ‘mock’ stimulation with L-15 media, or L-15 media containing 100 ng/ml IL10, IL6 or 20 μM ZPR-151. Donor and acceptor fluorescence intensity of individual cells within the bleached ROI was quantified before and after bleaching. Percentage FRET efficiency was calculated as %FRETeff = 100 x (D_post_ – D_pre_)/D_post_ where D_pre_ and D_post_ represent Clover-SHIP1 donor fluorescence intensity before and after bleaching, respectively.

### Immunofluorescence

Perimacs were seeded at 3 x 10^5^ cells per well in 18-well Ibidi μ-Slides (Ibidi GmbH, Martinsried, Germany) and allowed to adhere for 3 hours before washing with PBS to remove non-adherent cells. CD8+ T cells were seeded at 2 x 10^6^ cells in 12-well tissue culture plates. Cells were stimulated with either 100 ng/ml IL10 or 20 μM ZPR-151 for 2 or 20 minutes followed by 3 x PBS washes and fixing of cells in 4% paraformaldehyde for 15 minutes at room temperature. Cells were incubated with anti-mouse CD16/CD32 Fc Block (BD Pharmingen) for 1 hour followed by an overnight incubation at 4°C with either anti-SHIP1 antibody (P1C1) (Santa Cruz Biotechnology) or anti-STAT3 antibody (9D8) (ThermoFisher Scientific). Cells were then incubated with anti-mouse IgG (H+L)-Alexa Fluor 660 secondary antibody (ThermoFisher Scientific) for 1 hour, followed by, for perimacs, anti-CD11b-FITC antibody (BD Pharmingen) for 30 minutes. CD8+ T cells were mounted onto 18-well Ibidi μ-Slides prior to confocal microscopy and cells were stored in Ibidi Mounting Media supplemented with ProLong Gold antifade reagent with DAPI (Molecular Probes, Life Technologies). Cells were imaged on a Leica SP5II on DM6000 confocal microscope with a 63x/1.4-0.6 Oil PL APO objective using 405, 488 and 633 nm laser lines for excitation. Final images were scanned sequentially acquiring eight Z-stacks with a frame-average of four. Co-localization analysis was performed using ImageJ software by first combining individual z-stack confocal images then performing deconvolution and co-localization using CUDA deconvolution and JACoP plugins respectively. Pearson’s coefficient values were produced as a measurement of the degree of overlap between SHIP1 or STAT3 with CD11b (for Perimacs) or DAPI.

### Mouse Endotoxemia Model

Groups of 6-8 week old BALB/c SHIP1^+/+^ and SHIP1^-/-^ mice were intraperitoneally injected with either 1 or 5 mg/kg of LPS with or without co-administration of 1 mg/kg of IL10 or 5 mg/kg ZPR-MN100. Blood was drawn 1 hour later by cardiac puncture for determination of plasma cytokine levels by ELISA.

### Mouse Colitis Model

Colitis was induced in 6-8 week old BALB/c IL10^-/-^ mice by administering the colonic contents of conventional C57BL/6 mice diluted 1:10 in PBS by oral gavage. Mouse weights and fecal consistencies were monitored and colitis allowed to develop for 6 weeks. Ethanol (Vehicle) and ZPR-MN100 (3 mg/kg) was diluted in cage drinking water and dexamethasone (0.4 mg/kg) was administered every 2 days by oral gavage for 3 weeks. At the end of the dosing period, proximal, medial and distal colon sections were collected for paraffin embedding or stored in RNALater (Invitrogen, Mississauga, ON) for RNA extraction. Slides were prepared, stained with hematoxylin and eosin, and mounted by the UBC Department of Pathology and Laboratory Medicine Histochemistry Facility. Specimens were assigned pathological scores by two, independent, blinded investigators according to a method described by Madsen *et al* (Madsen et al., 2001). In brief, colonic inflammation was graded using a 4-point scale, scoring 0-3 for each of submucosal edema, immune cell infiltration, goblet cell ablation, and integrity of the epithelial layer. For analysis of mRNA expression, colon sections were homogenized and total RNA extracted as described above and analyzed by real time PCR using gene specific primers for IL17, CCL2, and GAPDH (normalization control).

### Expression of PAC2 for crystallography

LIC-HMT-PAC2 expression vector was transformed into *E. coli Rosetta(DE3) pLacI* cells. Overnight culture was inoculated with a 250-fold dilution to start the actual culture. The cells were grown at 37°C in LB medium (supplemented with 50 μg/ml of kanamycin and 34 μg/ml of chloramphenicol) with shaking at 225 rpm. When OD_600_ reached about 0.6, the culture was cooled down to room temperature before the addition of 0.4 mM isopropyl β-D-1-thiogalactopyranoside (IPTG) to induce the expression of recombinant protein. Cultures were left in the shaker overnight (usually 16-18 hours) at 22°C, and then collected by centrifugation (5000 g for 10 minutes at 4°C). The cell pellet was subsequently resuspended in lysis buffer (20 mM Tris-HCl pH 7.4, 350 mM NaCl, 10 mM TCEP, 5 mM imidazole, supplemented with 1X EDTA-free Protease Inhibitor Cocktail (PIC) (Roche Diagnostics, Laval, QC) and 25 μg/ml lysozyme), and lysed via sonication (2 cycles of 2 minutes pulse) on ice. Cell debris was removed by two rounds of centrifugation, first at 5000 g for 15 minutes at 4°C followed by 18000 rpm for 30 minutes at 4°C. Supernatant was filtered with a 0.45 μm filter and loaded onto a Talon Co^2+^-affinity column, previously equilibrated with Buffer A (20 mM Tris-HCl pH 7.4, 250 mM NaCl, 1 mM TCEP), and washed with 10 column volume (CV) of Buffer B (Buffer A + 5 mM imidazole). Bound proteins were eluted with 6 CV of Buffer C (Buffer A + 50 mM imidazole).

To remove the HMT tag, TEV protease (purified in house as a His_6_-tagged protein) was added to the eluted protein, which was then dialyzed against Buffer D (20 mM Tris-HCl pH 7.4, 250 mM NaCl, 1 mM TCEP) overnight at 4°C with gentle stirring. The dialyzed sample was loaded onto the Amylose column (New England Biolabs, Whitby, ON), and the flow through, which contained the untagged protein, was loaded onto the Talon column to remove the His_6_-TEV protease. The flow through from the Talon column was dialyzed against Buffer E (20 mM Tris-HCl pH 7.4, 25 mM NaCl, 1 mM TCEP) overnight at 4°C with gentle stirring, and then loaded onto the ResourceQ column (6 ml column volume) (GE Healthcare, Mississauga, ON), followed by washes with 3 CV of Buffer E. To elute the protein, Buffer F (20 mM Tris-HCl pH 7.4, 1000 mM NaCl, 1 mM TCEP) was used. A gradient from 25 mM NaCl (0% buffer G) to 200 mM NaCl (20% Buffer G) was used across 20 CV to separate the components in the protein sample. The fractions were analyzed by SDS-PAGE. PAC2 usually eluted from the ResourceQ column at ~130 mM NaCl. The purified protein was concentrated to about 5-10 mg/ml using Amicon concentrators with 30K MWCO (Millipore, Etobicoke, ON), and exchanged into the desired buffer. For protein crystallization, the desired buffer contained 50 mM Tris-HCl pH7.4, 25 mM NaCl and 0.5 mM TCEP. For Biolayer Interferometry, HMT-PAC2 proteins eluted from the first Talon column were directly purified on the ResourceQ column without cleavage of the HMT tag.

### Expression of PAC2-Avi tag for Biolayer interferometry

A sequence corresponding to Avi-tag (GLNDIFEAQKIEWHE) was added to the c-terminal end of the PAC2 in LIC-HMT-PAC2 expressing vector via standard restriction digestion and ligation. The LIC-HMT-PAC2-Avi expressing vector was then co-transformed into *E.coli BL21* cells with pBirAcm expression vector in 1:1 molar ratio. Overnight culture was inoculated with a 250-fold dilution to start the actual culture. The cells were grown at 37°C in LB medium (supplemented with 50 μg/mL of kanamycin and 10 μg/mL of chloramphenicol) with shaking speed of 225 rpm. When OD_600_ reached about 0.6, 5 mM of biotin in bicine buffer pH 8.3 was added to the culture to have final concentration of 125 μM of biotin. The culture was then cooled down to room temperature before the addition of 0.4 mM isopropyl-B-D-1-thiogalactopyranoside (IPTG) to induce the expression of the recombinant protein with Avi-tag. The rest of the method is identical as written in “Expression of PAC2 for crystallography”.

### Protein crystallization, data collection, phasing and refinement

Initial crystallization hits were obtained via sparse matrix screening in 96-well plates using commercially available crystallographic solutions (Qiagen, Toronto, ON). Optimization of crystallization conditions was performed in 24-well plate format using the hanging drop vapor diffusion method. Diffraction-quality protein crystals were obtained at 4-7 mg/ml protein at room temperature with 0.1 M HEPES-NaOH pH 6.7, 20% PEG1500 and 5 mM MgCl2. The PAC2-cc protein contained surface entropy reduction mutations (E770A, E772A, E773A) and aided in improving crystal quality. Unique fragments of crystal clusters of protein were soaked for 5 to 10 seconds in the crystallization solution containing 25% isopropanol, and flash-frozen in liquid nitrogen.

Diffraction data set were collected at the Advance Proton Source (APS) beamline 23-ID-D-GM/CA and processed with XDS through XDSGUI^45^. The phase problem was solved with an unpublished structure as search model in Phaser MR ^47^. The initial model was refined with COOT ^48^ and Refmac5 ^49^. Towards the final model occupancy refinement of sidechains was used in Phenix(Adams et al., 2010) and three TLS groups were defined. Data collection and refinement statistics are shown in **Table S1**. The model and data were deposited under protein database ID 6DLG.

### Small angle X-ray scattering

PAC1 samples in 190 μM in 50 mM TrisHCl (pH7.4) 150 mM NaCl, 1 mM TCEP and 2% EtOH with and without 570 μM ZPR-MN100 (6 fold molar excess). Dynamic Light Scattering (DLS) data for PAC1 and PAC1-ZPR-MN100 complexes were collected prior to SAXS data collection to confirm that all the samples are highly pure and suitable for data collection. The data collection was performed on a 3-pinhole camera (S-MAX3000; Rigaku Americas, The Woodlands, TX) equipped with a Rigaku MicroMaxþ002 microfocus sealed tube (Cu Kα radiation at 1.54 Å) and a Confocal Max-Flux (CMF) optics system operating at 40 W (Rigaku). Scattering data were recorded using a 200 mm multiwire two-dimensional detector. The data for both samples and buffer were collected for 3 h for each sample within the range of 0.008 ≤ s ≤ 0.26 Å-1 and processed according to the method previously described, where s = 4πsin θ/λ (Patel et al., 2011, Patel et al., 2010, Patel et al., 2012). The Normalized Spatial Discrepancy (NSD) of the non-liganded and liganded PAC1 models were 0.6 and 1.0 respectively.

### Biolayer interferometry

The binding affinity between the PAC2 protein and small molecule allosteric regulators was examined via bio-layer interferometry (BLI) experiments using super-streptavidin (SSA) biosensor tips and an Octet Red 96 instrument (ForteBio, Fremont, CA). SSA biosensor tips were hydrated in assay buffer 20 mM Tris-HCl (pH 7.4), 150 mM NaCl, 10 mM MgCl2, 0.5 mM TCEP, 0.2% Tween-20 prior to protein immobilization. 0.5 ug/mL of protein was immobilized to the SSA biosensor overnight at 4°C. After immobilization of protein to the biosensor, the tips were blocked with 0.1% BSA for 90 minutes followed by 20 minutes of wash with assay buffer supplemented with 1% EtOH. The kinetic measurement was done at 30°C with orbital flow of 1,000 RPM. The baseline was achieved with the assay buffer supplemented with 1% EtOH for 60 s. The association was measured for 600 s at an analyte concentration of 20 μM followed by dissociation for 300 s in the same buffer as the baseline. The raw data was analyzed using the Octet Red Data Analysis software (ver. 8.2). The raw data were aligned to the baseline and subtracted using both single and double reference subtraction.

## QUANTIFICATION AND STATISTICAL ANALYSIS

The band quantification of all immunoblots were performed using LI-COR Odyssey imaging system and Image Studio^™^ Lite software (LI-COR Biosciences, Lincoln, NE). GraphPad Prism 6 (GraphPad Software Inc., La Jolla, CA) was used to perform all statistical analyses. Statistical details can be found in figure legends. Values are presented as means ± standard deviations. Unpaired *t* tests were used where appropriate to generate two-tailed P values. One-way or Two-way ANOVA were performed where required with appropriate multiple comparisons tests. Differences were considered significant when *p*≤0.05.

## DATA AND SOFTWARE AVAILABILITY

The X-ray crystallography data that supports the findings of this study has been deposited in the Worldwide Protein Data Bank (wwPDB) under the ID code, PDB ID 6DLG.

