## Supplemental Figures for "Interleukin-10 and Small Molecule SHIP1 Allosteric Regulators Trigger Anti-Inflammatory Effects Through SHIP1/STAT3 Complexes"

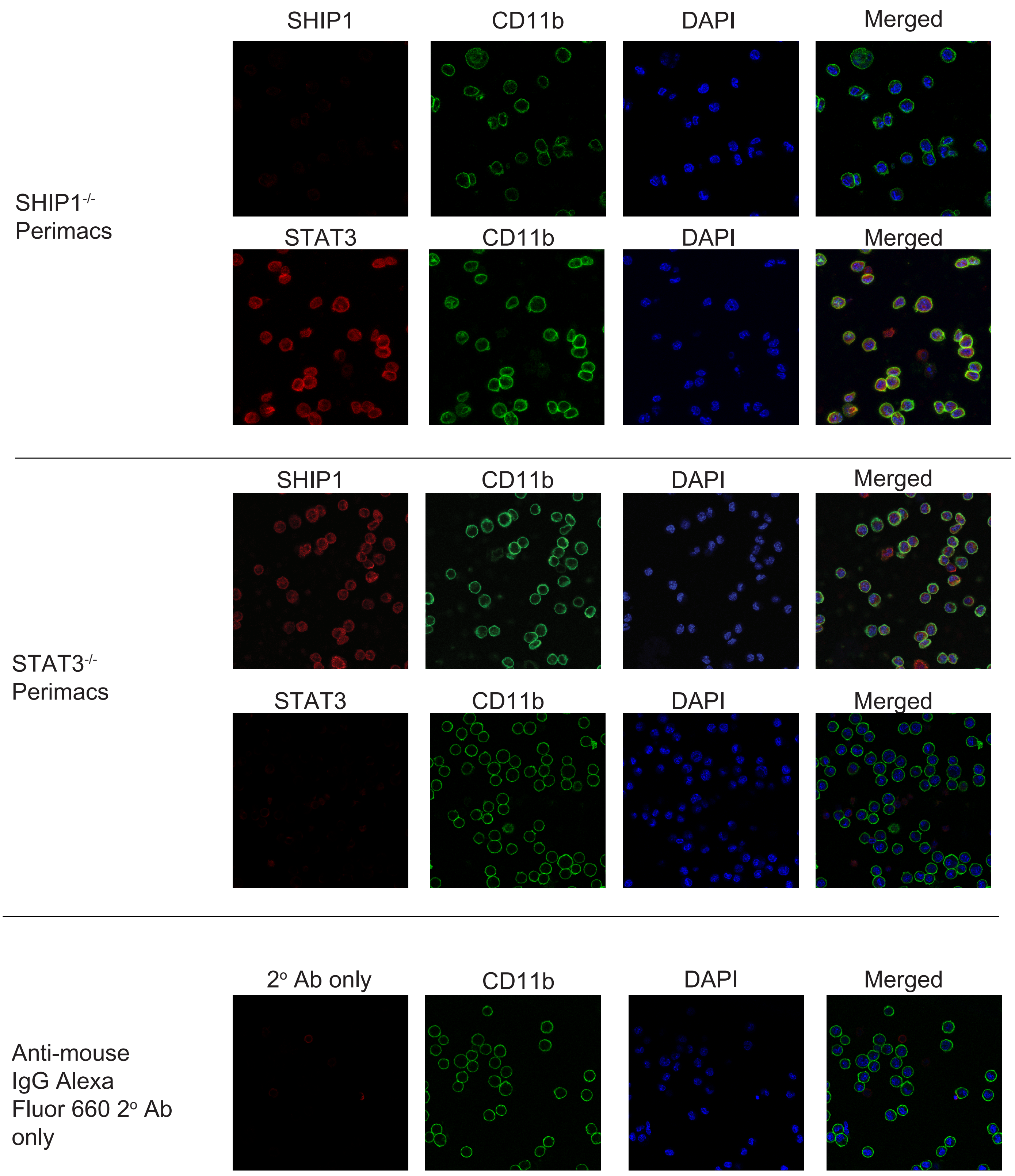


**Figure S1,**

**Related to Figure 4. SHIP1 and STAT3 immunofluorescence staining controls**

Representative confocal microscope images of unstimulated SHIP1$^{-/-}$ and STAT3$^{-/-}$ perimacs stained with anti-SHIP1 and anti-STAT3, CD11b antibodies and DAPI. Representative images of perimacs stained with anti-mouse IgG Alexa-Fluor 660 secondary antibody without anti-SHIP1/STAT3 primary antibody staining.

### A K681A PAC2

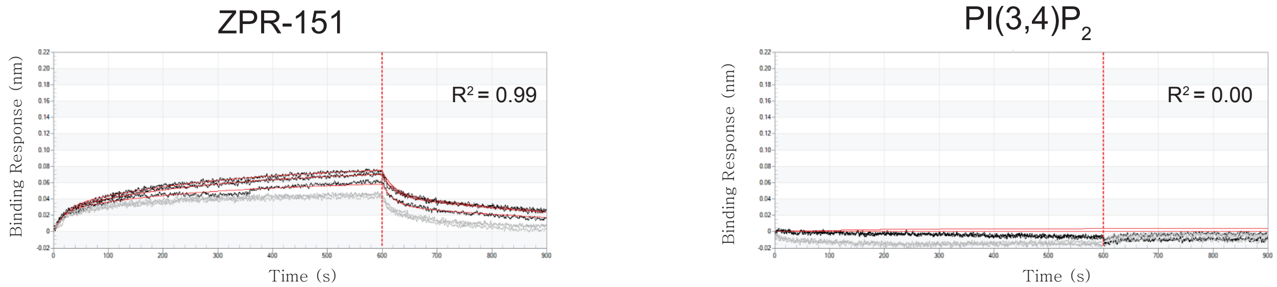


### B WT PAC2

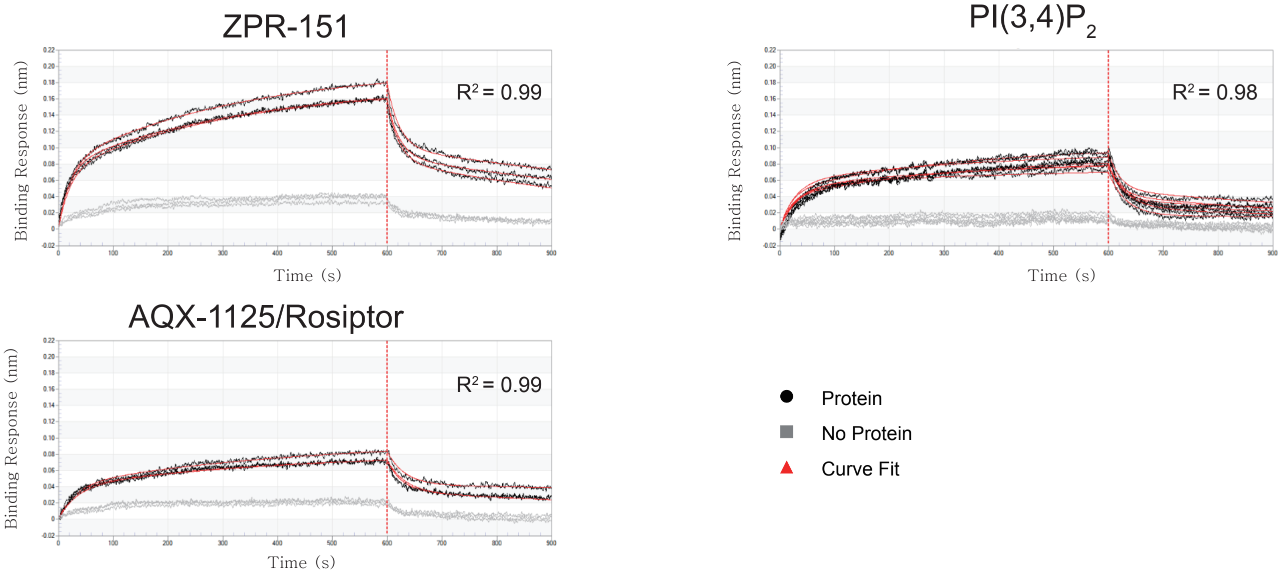


# C

| Protein | Compound | Replicates | $k_{on}$ ($M^{-1}s^{-1}$) | | $k_{off}$ ($s^{-1}$) | |
|---|---|---|---|---|---|---|
| | | | Mean | Std. Error | Mean | Std. Error |
| WT | ZPR-151 | 3 | 147 | 46.2 | 0.062 | 0.000749 |
| WT | $PI(3,4)P_2$ | 3 | 42.6 | 21.9 | 0.0288 | 0.000435 |
| WT | AQX-1125 | 3 | 65.5 | 53.4 | 0.0485 | 0.000943 |
| K681A | ZPR-151 | 3 | 69.7 | 65.2 | 0.0477 | 0.00116 |
| K681A | $PI(3,4)P_2$ | 3 | 250 | 1.163 | <1.0E-07 | |

#### Figure S2

**Related to Figures 6 and 7. Bio-layer interferometry (BLI) curves from WT and K681A PAC2 loaded BLI sensors exposed to 20 mM of ZPR-151, $PI(3,4)P_2$ or AQX-1125/Rosiptor**. **(A)** Representative BLI curves for binding of 20 µM ZPR-151 and $PI(3,4)P_2$ to K681A PAC2. **(B)** Representative BLI curves for binding of 20 µM ZPR-151, AQX-1125 and $PI(3,4)P_2$ to wild-type (WT) PAC2. The black and gray lines (n=3) show the observed binding curves to PAC2 and no protein loaded biosensors respectively. The red lines show the curve fit of each black line. The curves are fitted to 2:1 heterogeneous ligand model indicating possibility the protein is heterogeneous (FortéBio. Technical Note 16: Small Molecule Binding Kinetics. https://www.fortebio.com/literature.html) **(C)** Representative of $k_{on}$ and $k_{off}$ values obtained from the fit curves.

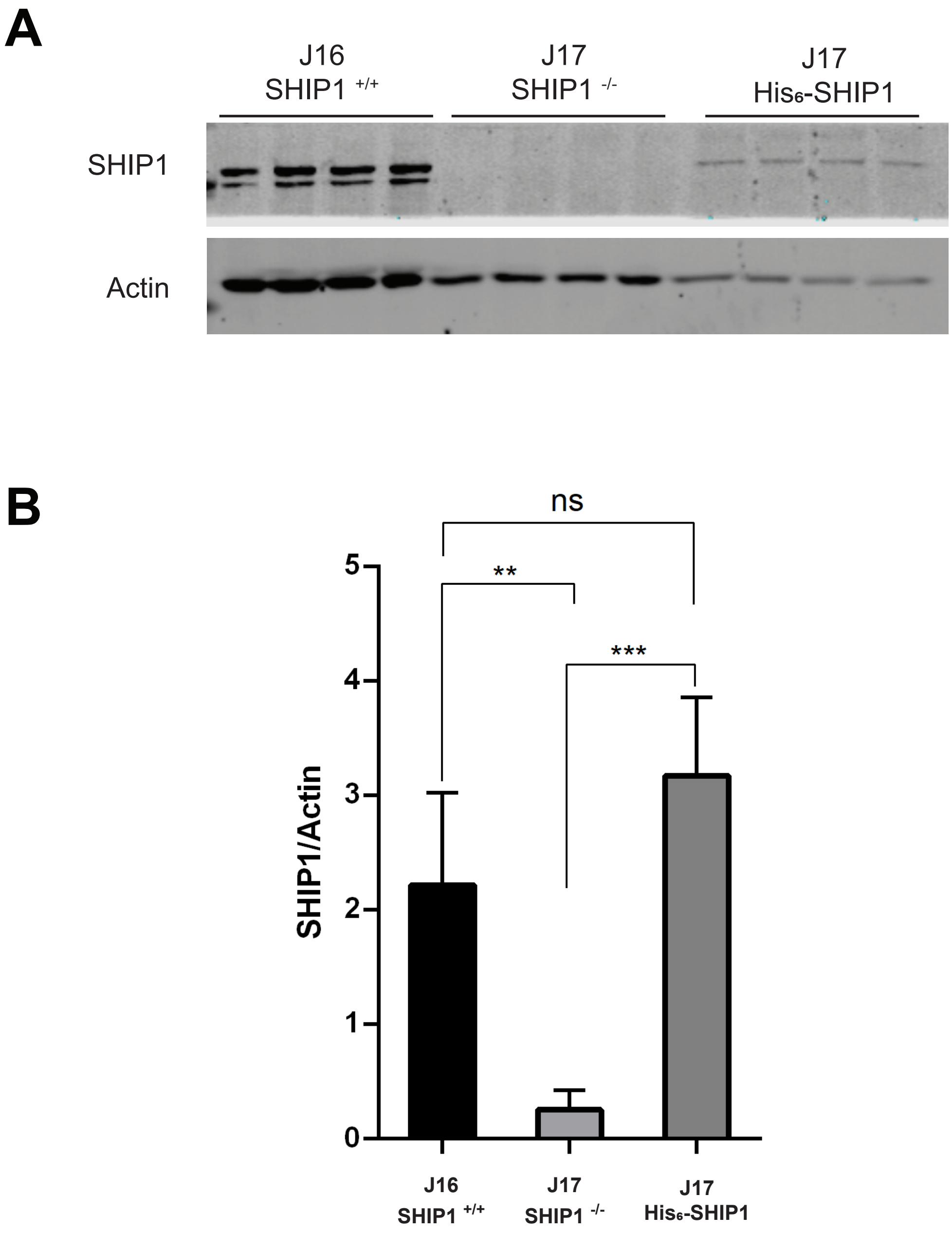


### Figure S3

**Relative Expression of endogenous wild-type SHIP1 and the transduced HIS-SHIP1 proteins.** Lysates from J16 SHIP1 $^{+/+}$, J17 SHIP1 $^{-/-}$ and J17 SHIP1 $^{-/-}$ cells reconstituted with His$_6$-SHIP1 reconstituted were collected. (A) Expression of endogenous SHIP1 and transduced His-SHIP1 level was detected by immunoblotting. (B) Quantification of SHIP1 protein level normalized to actin. (One Way ANOVA, *** $p < 0.001$, ** $p < 0.01$, ns = not significant).

| | |
|---|---|
| Protein sample<br>PDB ID | SHIP1 PAC2-cc<br>6DLG |
| **Data collection** | APS 23ID-D |
| Wavelength (Å) | 1.03319 |
| Space group | P 21 21 21 |
| Unit cell parameters (Å) | a=45.10, b=73.20, c=124.21 |
| Unit cell angles (°) | α=90.0, β= 90.0 γ= 90.0 |
| Resolution (Å)* | 47.36-1.50 (1.59-1.50) |
| $R_{meas}$ (%) | 5.9 (78.7) |
| $CC_{1/2}$ (%) | 99.9 (68.2) |
| $I$ / σ($I$) | 13.79 (1.85) |
| Completeness (%) | 98.9 (98.6) |
| No. unique reflections | 66070 (10528) |
| Redundancy | 3.28 (3.29) |
| Wilson B factor (Å$^2$) | 25.38 |
| **Refinement** | Phenix 1.13-2998 |
| Resolution (Å) | 38.4-1.50 |
| Solvent content (%) | 38 |
| $R_{work}$ / $R_{free}$ (%) | 17.85/20.04 |
| Ramachandran plot | |
| favoured / outlier (%) | 99.3 / 0.2 |
| No. of atoms | |
| Protein | 3628 |
| Isopropanol | 4 |
| Water | 246 |
| $B$-factors (Å$^2$) | |
| Protein | 24 |
| Isopropanol | 26.6 |
| Water | 30.9 |
| RMSD bond lengths (Å) | 0.008 |
| RMSD bond angles (°) | 0.966 |

*Values of the highest resolution shell are listed within parentheses

### Table S1

**Related to Figure 6.** Protein crystallization data collection and refinement statistics
